# Loading causes molecular damage in fibrin fibers

**DOI:** 10.1101/2025.05.08.652948

**Authors:** Sajjad Norouzi, Matthew J. Lohr, Mayar T. Ibrahim, Christian M. Jennings, Daniel Wang, Pengyu Ren, Manuel K. Rausch, Sapun H. Parekh

## Abstract

Blood clotting is the body’s natural reaction in wound healing and is also the cause of many pathologies. Fibrin – the main protein in the clotting process provides clots’ mechanical strength by forming a scaffold of complex fibrin fibers. Fibrin fibers exhibit high extensibility and primarily elastic properties under static loading, which differ from in vivo dynamic forces. In many biological materials, the mechanical response changes under repeated loading/unloading (cyclic loading). Using lateral force microscopy, we show fibrin fibers possess viscoelastic behavior and experience irreversible damage under cyclic loading. Cross-linking results in a more rigid structure with permanent damage occurring mostly at larger strains, which is corroborated by computational modeling of fibrin extension using a hyperelastic model. Molecular spectroscopy analysis with broadband coherent anti-Stokes Raman scattering spectroscopy in addition to molecular dynamic simulations allow identification of the source of damage, the unfolding pattern, and inter and intramolecular changes in fibrin. The results show partial recovery of protein’s secondary and tertiary structures under load, providing deeper understanding of fibrin’s unique behavior in wound healing or pathologies like stroke and embolism.

## Introduction

Fibrin, the pivotal protein in blood clots, plays a crucial role in wound healing and thrombosis as a scaffold that unites clot components to prevent bleeding and promote tissue repair ^1,2^. The hierarchical structure of fibrin scaffolds arises from the polymerization of its precursor, fibrinogen, into fibrin fibers. Fibrinogen comprises disulfide-bonded pairs of Aα, Bβ, and γ chains ^3^. Together, these chains form two distal (D-domain) and one central (E-domain) globular domains, with each D- and E-domain connected by a three-stranded, α-helical coiled-coil structure. Through the coagulation cascade, thrombin cleaves fibrinopeptide A and B from Aα and Bβ chains of fibrinogen, leaving α and β chains and creating knobs ‘A’ and ‘B’ and holes ‘a’ and ‘b’ as fibrin molecules ^3,4^. These molecules grow longitudinally through knob-hole interactions, forming long 2-strand thick protofibrils that can aggregate laterally to create fibrin fibers. Protofibrils are further stabilized through the αC domain – the C-terminal part of the α chain that enhances the lateral aggregation of protofibrils ^5,6^. Finally, Factor XIII (FXIII) further stabilizes the clot by forming covalent bonds in αC domain and the C-terminal part of the γ chain known as γ-nodule ^7^.

Blood clots naturally experience mechanical loads in the body. They feel shear from blood flow and tensile forces from platelet contraction and muscle movements. The mechanical properties of these clots are crucial for withstanding forces, facilitating wound closure, and recommending treatment strategies for their effective removal in conditions such as thrombosis and stroke ^8^. Various techniques, from atomic force microscopy (AFM) to dynamical mechanical analysis and mechanical rheology, have been employed to investigate clot mechanical features at different scales, spanning from a single fibrinogen molecule to the entire clot ^9–12^. Regardless of the scale, clots or fibrin gels exhibit high extensibility and viscoelastic properties with stress-stiffening behavior; that is, an increased elastic modulus with strain ^13–16^.

Single molecule spectroscopy and molecular dynamic (MD) simulations have demonstrated fibrinogen’s unfolding under force, starting with the unfolding in the γ-nodule and helical coiled-coil and propagating through other regions ^17^. These studies show how molecular rearrangements within fibrinogen accommodate external forces. Interstingly, while the αC region shows limited contribution to the single molecule (fibrinogen) mechanical properties, this region does influence single fibrin fiber and network mechanical properties ^17–19^. Cross-linking by FXIII in both αC region and γ-nodule also alters fibrin’s stiffness and extensibility ^18,19^. Notably, these insights have primarily relied on static (singular) loading under tension, compression, or shear, which do not exactly align with in vivo dynamic forces, such as those applied during the respiratory cycle or muscle movements. Many biological materials exhibit different mechanical behavior under cyclic loadings compared to common acyclic scenarios ^20–22^. In cyclic loadings, a material undergoes repeated loading/unloading over time, such as with peristaltic forces that occur during respiration. While techniques like AFM or optical tweezers provide insights into fiber-scale behavior ^9,23^, they fall short in detailing how force propagates within the complex fibrin molecule and which parts of the molecule respond to the external force. Spectroscopic techniques, such as Fourier transform infrared (FTIR) and broadband coherent Raman anti-Stokes scattering (BCARS), offer structural information at the molecular level ^24,25^. FTIR and BCARS have revealed the change in the protein secondary structure through α-helix to β-sheet transition under tension or compression, which was attributed to the unfolding of fibrin’s helical coiled-coil domain. Our previous study demonstrated the involvement of the D-domain with β-sheet structures in fibrin’s response to the external force ^26^, consistent with β-strands unfolding in MD simulations of the fibrinogen molecule^17^. However, the role of unfolding in this domain and its interaction with the coiled-coil domain is unclear. Which domain unfolds first and to what extent? Answering this question may help clarify what parts of the molecule are responsible for the observed inconsistent increasing/decreasing trend of the α-helix to β-sheet ratio under increasing shear strain ^26^. Additionally, it remains uncertain whether force can induce any permanent molecular-level damage in fibrin and the source of that damage, as similar to many biological materials, the loading history may change the subsequent fibrin’s response to the load in a cyclic loading scenario ^20,21,27^.

In this study, we employ lateral force microscopy as an application of AFM to explore the mechanical response of single fibrin fibers in different cross-linking scenarios and under cyclic loadings ^28^. Subsequently, we use BCARS spectroscopy and MD simulations to elucidate fibrin’s molecular behavior under cyclic loading to trace damage propagation in different molecule parts. Our observations indicate that increased cross-linking by FXIII enhances fibrin’s rigidity, reducing permanent damage to fibers under tensile loading. Conversely, uncross-linked fibrin fibers show irreversible damage even at small strains, which intensifies as strain increases. BCARS data consistently depict changes in the protein’s secondary and tertiary (3-dimensional (3D) conformation) structure under force for uncross-linked fibers. However, this behavior is more vivid for cross-linked fibers at larger strains. Intermolecular interactions, particularly in cross-linked fibers, change under load, which suggests that protofibrils’ lateral configuration differs before versus after loading. This study provides insights into fibrin’s response to cyclic loading, deepening our understanding of in vivo blood clot behavior since the material fundamentally changes with loading history.

## Materials and Methods

### Sample preparation

To create individual fibrin fibers for AFM experiments, fibrinogen (FIB1, Enzyme Research Laboratories) was dissolved in FB1x buffer (20 mM HEPES (BP310-100, Fisher Scientific) and 150 mM NaCl (S271-3, Fisher Scientific) at pH 7.4) to prepare a 10 mg/mL stock solution. For fluorescence imaging, unlabeled fibrinogen was mixed with Alexa 488-labeled fibrinogen (F13191, Thermo Fisher) at a 9:1 molar ratio. Fibrinogen and human alpha thrombin (HT 1002a, Enzyme Research Laboratories) were then combined with FB1x buffer containing 5 mM CaCl_2_ (746495, Sigma Aldrich) to achieve final concentrations of 1 mg/mL fibrinogen and 0.1 U/mL thrombin. FIB1 consists of some FXIII, resulting in partially cross-linked fibrin fibers. Next, we applied 10 μL of the resulting solution onto a ridged substrate, fabricated as outlined in ^9^. Briefly, two drops of optical adhesives (NOA 60, Norland Optical Adhesives) were cured by ultraviolet light between a mold made of polydimethylsiloxane (PDMS) (Research micro stamps) with 30 μm wide trenches and ridges and a cover glass. The optically cured substrate was plasma-cleaned for hydrophilicity before pouring the fibrin gel. After incubating the fibrinogen for one hour at 37°C in ∼ 90% humidity, it was rinsed with FB1x buffer to remove excessive salts and weakly bound fibers. Throughout the entire experiment, the sample remained hydrated in FB1x buffer.

To inhibit the cross-linking in FIB1, we used 1,3-Dimethyl-4,5-diphenyl-2-[(2-oxopropyl) thio] imidazolium, trifluorosulfonic acid salt (D004) (Zedira). D004 was dissolved in dimethyl sulfoxide (DMSO) to make a 20 mM solution, as described in ^29^. Then, 2 μL of this solution was added (400 μM final concentration) to the fibrinogen solution before adding thrombin.

For fully cross-linking fibrin fibers, we used exogenous Factor XIII (HFXIII 1313, Enzyme Research Laboratories), with a final concentration of 8 U/mL ^24^. It was added to thrombin solution in the presence of CaCl_2_ for full cleavage and then mixed with fibrinogen solution.

In the BCARS experiment, we created fibrin gels with higher fibrinogen and thrombin concentrations to increase Raman scattering signal (final fibrinogen and thrombin concentrations were 7 mg/mL and 1 U/mL, respectively). The fibrin gel was formed by the fibrinogen/thrombin solution bridging two 24 mm x 60 mm plasma-cleaned coverslips (48393-106, VWR) with a 2-3 mm gap on a manual tensile device. The gel solution was supported by a third square 22 mm x 22 mm cover glass (48366-227, VWR) coated with PEG (6-9) silane to prevent protein adhesion^30^. The lateral boundaries on two coverslips were established with a hydrophobic marker. Subsequently, the gel was incubated for an hour at 37°C with ∼90% humidity. Then, the bottom PEG-silanized cover glass was gently detached and replaced with another square cover glass spaced ∼ 175 μm from the gel to hold the buffer and hydrate the gel.

### Lateral force microscopy

Fibrin mechanical testing was conducted using the MFP-3D atomic force microscopy (Asylum Research) equipped with inverted epifluorescence microscopy (Olympus IX 70). Optical images were acquired with a 30X, 1.05 NA silicone oil immersion objective (UPLSAPO30XS, Olympus). To ensure accurate mechanical measurements with enough resolution, soft cantilevers with nominal normal spring constants of 200 pN/nm and 20 pN/nm (PPP-LFMR, qp-SCONT, NanoAndMore) were used. All cantilevers were calibrated for lateral force microscopy (LFM) using the geometrical features and spring constant in normal mode ^31^. Subsequently, when the cantilever tip is approximately in the fiber’s axial plane, the cantilever is moved laterally, impinging on a single fibrin fiber and starting the pulling process ^9^. The cantilever speed was set to 1 μm/s, as no differences were observed in the mechanical responses of fibrin fibers at varying loading rates (**Fig. S1**). In addition to recording force based on the calibrated lateral AFM signal and stage cantilever displacement, the fiber is imaged for length measurements and strain calculations. Strain is calculated as the ratio of change in fiber’s length to the initial length.

For force relaxation tests, the fiber is pulled to a certain strain (1 μm/s is the cantilever lateral velocity) and held there while recording the force until being pulled to a larger strain. In cyclic loading experiments, the fiber was pulled to a certain strain twice to differentiate viscoelastic energy dissipation and permanent damage.

### Broadband coherent anti-Stokes Raman scattering (BCARS) microscopy

For molecular microscopy, we used the BCARS hyperspectral microscope that was previously used for fibrin measurements. Briefly, a laser producing sub-100 picosecond pulses at 1 MHz repetition rate at 1064 nm and a broadband supercontinuum with a spectrum ranging from 1100 nm to 2400 nm (Leukos Opera HP, Leukos) was used for excitation. The two beams were focused by a 100X, 0.85 NA objective (LCPLN100XIR, Olympus). The signal was collected in transmission by a 20X, 0.4 NA objective (M-20X, MKS Newport).

The formed fibrin gel on the tensiometer was mounted on the BCARS stage. The gel was fixed at one end while the other end could move by a manual micrometer. The uniaxial tension was applied by turning the micrometer, and the strain was calculated based on the ratio of deformation to the initial gauge length. In the cyclic loading, the gel was pulled to each strain twice and then returned to the relaxed state. The pixel size was chosen 1μm x 1μm, and the CCD integration time for each spectrum was ∼ 200 ms to maximize the signal while avoiding saturation. The experiment was conducted for both uncross-linked and partially cross-linked (without inhibiting or adding cross-linkers to FIB1) fibrin gels with four and five replicas, respectively.

### BCARS signal processing

BCARS raw signals were converted into Raman spectra based on the methodology outlined in literature ^4,5^. BCARS spectra with an average spectral resolution of 4 cm^−1^ was recorded over a range of 700 cm^−1^ to 4000 cm^−1^ Raman shift. After phase retrieval, all the spectra were normalized based on the total Amid I area (1600 cm^−1^ to 1700 cm^−1^), showing the protein secondary structure. Subsequently, the background was subtracted from each spectrum based on the intensity at the smallest Raman shift. Each spectrum was at least the average of 3 x 3 pixels. For molecular analysis, the ratio of α-helix (mean intensity in the range of 1645 cm^−1^ to 1655 cm^−1^) to β-sheet (mean intensity in the range of 1669 cm^−1^ to 1679 cm^−1^) ^32^, the intensity at 3065 cm^−1^ (mean intensity in the range of 3060 cm^−1^ to 3070 cm^−1^) as the indication of aromatic amino acids ^33^, and the intensity at 2934 cm^−1^ (the mean intensity in the range of 2929 cm^−1^ to 2939 cm^−1^) as CH_3_ symmetric vibrations ^32^ were plotted for each strain in violin plots ^34^. Statistical analyses were done based on non-parametric Wilcoxon signed-rank test.

### Computational modeling of cyclic loading

We developed a computational model to capture the visco-hyperelastic and damage properties of the fibrin fibers under cyclic loading. Specifically, we fit a quasi-linear visco-hyperelastic damage model to data on force, time, and displacement from the cyclic loading tests using custom MATLAB scripts. We modeled fiber deformation as a combination of two uniaxial extension problems and assumed that its deformation was isochoric. Note that, because we did not have accurate measures of diameter, we used a diameter of 150 nm for all samples. This was chosen as it is within the diameters reported for individual fibrin fibers by Belcher et al. ^35^. Using this fixed value allowed us to compare the parameters across samples; however, it did not allow us to make accurate comparisons of the fibers’ stiffnesses. Also, note that we do not include the bending of the fiber in our model; this is because the bending does not contribute significant energy to the system at the strains in these mechanical tests. Our supplemental document contains a figure showing the distribution of energies between bending and stretching (**Fig. S2**). Using the initial position of the AFM cantilever along the fiber and its horizontal and vertical displacements, we calculated the stretch that the fiber would feel due to the moving beam. To model the visco-hyperelastic properties of the fibers, we implemented a quasilinear viscoelastic model from Simo and Hughes ^36^. In this model, the principal components of the visco-hyperelastic stress take the form

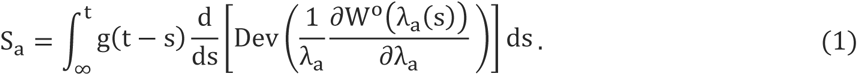

Here, a indexes the principal coordinate axes and 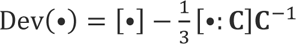. Here, **C** is the right Cauchy-Green tensor and is defined as 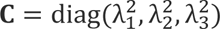. Note that boldface denotes that a given variable is a second-order tensor. For our hyperelastic strain energy function, W°, we used a one-term Ogden model, which is an isotropic model defined as

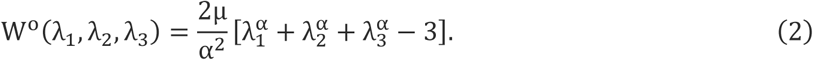

The Ogden model uses the principal stretches, λ_i_ (i = 1,2,3), of the deformation. Because we did not have equipment capable of measuring the volume change of the fiber, we assumed that there was no volume change. Thus, we excluded the volumetric portion of the stress from Equation 1. Due to our incompressibility assumption, the stretches of our uniaxial loading are found to be λ_1_ = λ and λ_2_ = λ_3_ = 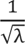. Here, λ is the stretch in the direction of the loading, x_1_. The parameter α governs the strain-stiffening behavior of the material and μ is the shear modulus of the material. Note that W° is the isochoric portion of the strain energy.

We used a two-term (N = 2) Prony series for our normalized relaxation function, g(t). This modulus, which captures the time-dependent behavior of the fiber, is defined as

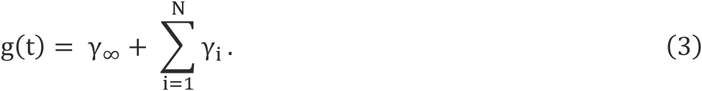

Here, τ_i_ > 0 are the relaxation time constants and γ_i_ ∈ [0,1] are moduli subject to the following constraint

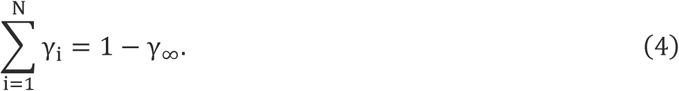

Here, γ_∞_ is the long-term modulus corresponding to a fully relaxed material.

We model the progressive, Mullins-like damage of fibrin fibers through the Simo damage model^37^. In this model, the damage is a function of the maximum stretch a body has experienced in its loading history. The body only undergoes damage when it is stretched beyond its previous maximum stretch. To this end, our effective stress, **S̅**, is defined as

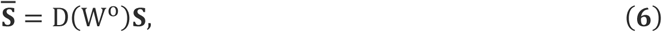

where D(W°) ∈ [0,1] is a damage function with the form

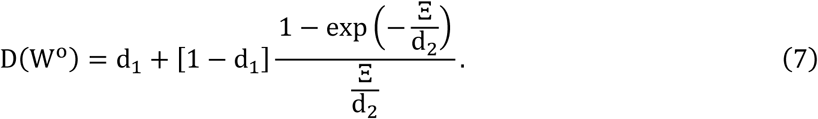

Here, d_1_ ∈ [0,1] and d_2_ > 0 are parameters governing the damage evolution, and Ξ is defined as

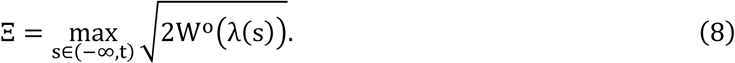

Note that this damage formula captures the Mullins-like effect seen in repeated loading of fibrin fibers ^38^. We implemented the integration algorithm for nonlinear viscoelasticity from Simo and Hughes ^36^ to calculate S_a_ in Equation 1. Also, note that we fit the model to the lateral force using data on force, time, and displacement. Our error function for this optimization process was

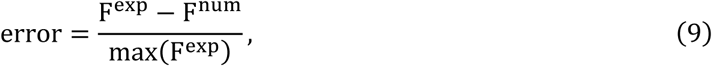

where F^exp^ is the force measured in the experiment, and F^num^ is the force computed by the numerical model. We minimized the total error using a nonlinear least squares solver with the trust-region-reflective algorithm within MATLAB. Note that we fit all model parameters (μ, α, γ_i_, τ_i_, d_1_, d_2_) at once.

### Molecular dynamics simulations of single molecule fibrinogen stretching

The crystal structure of a single fibrinogen molecule (PDB ID: 3GHG) was obtained from the Protein Data Bank. System preparation was conducted using the GROMACS package ^39,40^ with the CHARMM27 force field ^41^. The default protocol was followed, including solvation in a water box (cutoff: 10.0 Å) using the TIP3 water model, and neutralization to physiological salt concentration by adding Na⁺ ions. The system was then energy-minimized and equilibrated under canonical ensemble with constant Number of particles, Volume, and Temperature (NVT) and isothermal-isobaric ensemble with constant Number of particles, Pressure, and Temperature (NPT) conditions. Next, the equilibrated system was subjected to two independent pulling simulations using CustomBondForce in OpenMM 8 package 42. The pulling force was applied to ILE394 in γ-nodule 1 at one end of the fibrinogen molecule, while GLY395 in γ-nodule 2 at the other end of the protein remained fixed. To achieve the corresponding experimental strain of 16.67%, a pulling force constant of 1000.0 kJ/molꞏnm was applied (Pull 1, **Fig. S3**), followed by an 8 ns relaxation phase (Relax 1, **Fig. S3**) without external force to reach the lowest possible strain. For the second pulling simulation (Pull 2, **Fig. S3**), a lower force constant of 50 kJ/molꞏnm was used to achieve the same strain, followed by a 5 ns relaxation phase (Relax 2, **Fig. S3**). The strain was calculated as the ratio of the molecule extension to the fibrinogen initial length between two ends of the protein (ILE394 & GLY395).

During the initial pulling simulation (Pull 1), the target strain of 16.67% was reached within 200 ps and remained stable for the remainder of the 1 ns pulling time (**Fig. S3**). The final frames from two independent Pull 1 replicas were selected as representative structures (**Fig. S3**) and used as the starting state for the subsequent Relax 1 phase. Relax 1 was run for 8 ns, but after 6 ns, strain spikes were observed in both replicas (**Fig. S3**). Therefore, the final frame at 6 ns was chosen as the representative structure for Relax 1 and used as the seed structure for the next pulling simulations. For the second pulling simulation (Pull 2), a lower pulling force constant (k = 50.0 kJ/molꞏnm) was applied. The 16.67% strain was achieved at 400 ps for Replica 1 and 800 ps for Replica 2, with stable strain values reflecting structural stability (**Fig. S3**). The final frames were selected as representative structures. These selected structures from Pull 2 were used to initiate the Relax 2 phase, which was run for 5 ns in two independent replicas. The last frames of these simulations were chosen as representative structures for this phase.

Ramchandran plots were generated using MDAnalysis2.9.0 ^43,44^. The α-helix/β-sheet ratio was determined based on the number of points within the areas assigned to each secondary structure. α-helices were identified within ϕ angles ranging from −120° to −30° and 60° to 90°, while ψ angles ranged from −60° to −30° and 0° to 60°. β-sheets were also characterized by ϕ angles between - 180° and −45° and ψ angles from 60° to 180°.

## Results

### Single fibrin fiber mechanics show stress relaxation

Using the AFM for LFM, we deformed single fibrin fibers inspired by work by Guthold and colleagues ^9,28^. We initially conducted force relaxation experiments for single fibrin fibers that are: uncross-linked, partially cross-linked, and fully cross-linked by FXIII. Single fibers were pulled to a specific strain and held at that strain while recording the force signal. The force signals for different strain steps are shown in **Figure 1A-C**, with the force recorded on the left axis and the corresponding strain steps on the right axis. Notably, as the strain increases, fibrin fibers, independent of the cross-linking state, relax more. This behavior initiates at much lower strains for uncross-linked fibers (less than 5% strain), indicating greater energy dissipation and, hence, more viscoelastic behavior. In cases where the material is purely elastic, there is no stress relaxation, and the force remains constant, as observed in the fibrin fibers subjected to very small strains. To assess and compare the effect of cross-linking on viscoelastic properties, we calculated the mean square error (MSE) for the force response at each strain step. The MSE value is the deviation of the data from the mean force value, a representative constant force, as would be the case for fully elastic materials. **Figure 1D** shows the MSE for different groups at varying strains, highlighting how uncross-linked fibers deviate from the mean value (the ideal elastic response) even at very small strains (5% strain). Conversely, cross-linking results in more elastic fibers, delaying energy dissipation until larger strains are reached.

**Figure 1.**
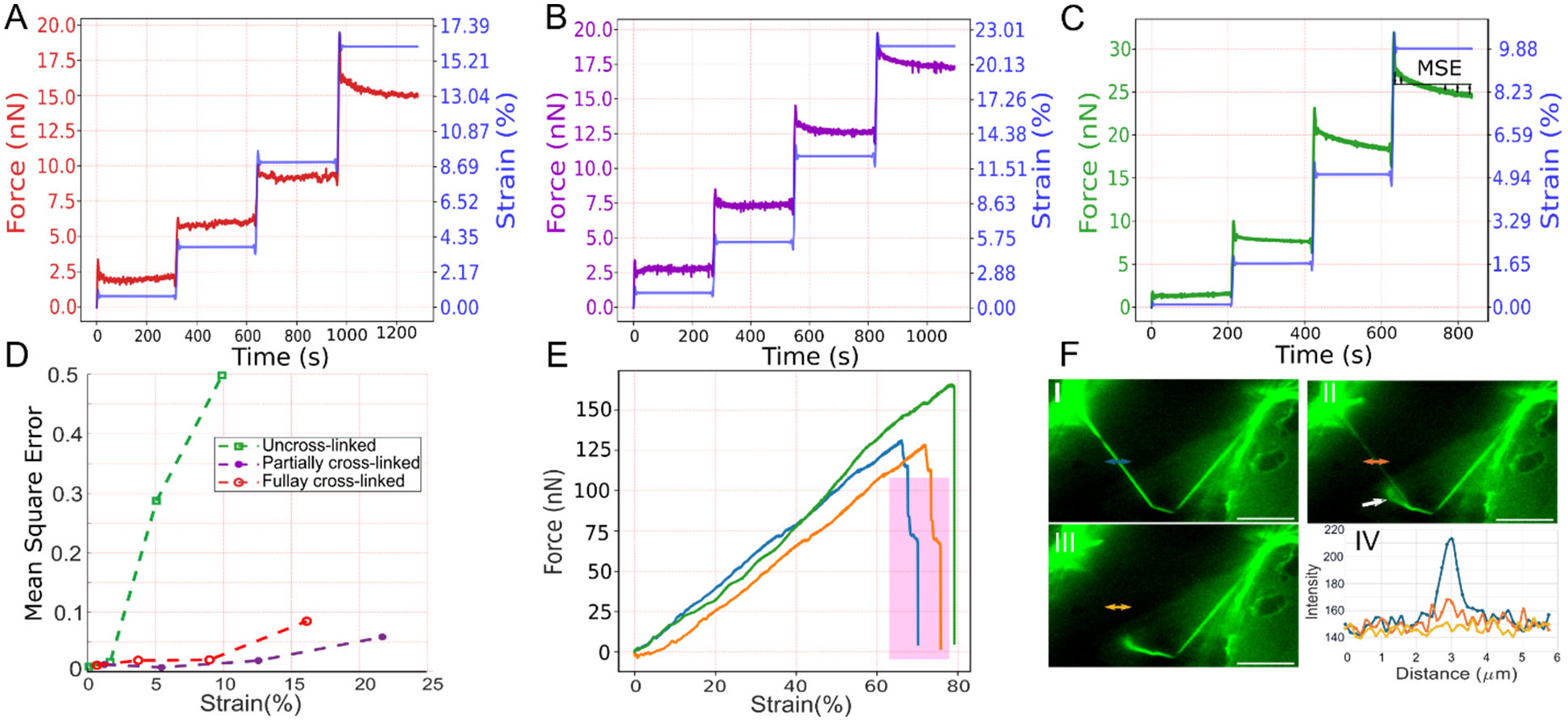
Relaxation tests for fully cross-linked (A), partially cross-linked (B), and uncross-liked (C) fibrin fibers. (D) Mean square error (MSE) of the force in each strain step for three groups as an indication of how much the relaxation curve deviates from the mean force value. (E) Force-strain curve for three different partially cross-linked fibrin fibers. The shaded area shows the region where catastrophic breakage of fibers often occurred. (F) The fraying pattern of a fully cross-linked single fibrin fiber at the breakage point. I to III are three consecutive snapshots of fiber rupture. The 2-sided arrows’ intensity is plotted in IV and the white arrow shows the frayed fiber. The scale bar is 10 μm.

In **Figure 1E**, the force-strain response of three partially cross-linked fibrin fibers is presented until rupture. Notably, instead of stress, we plot force, considering the small scale of fibrin fibers, where the continuum mechanics assumption of a uniform rod can be inappropriate. Previous work showing that, as fiber diameter increases, fiber’s elastic modulus decreases ^9,45,46^, supports the idea that fibrin fibers may not be uniform rods. Thus, calculating stress by normalizing force with the fiber cross-sectional area may not properly account for the size-dependent heterogeneous properties in each cross-section. The highlighted region in **Figure 1E** shows the force dropping in two or three steps before full rupture. **Figure 1F**(I and II) provides two consecutive snapshots of a fully cross-linked fibrin fiber just before rupture in **Figure 1F**(III). The intensity of the image through a cross-section is plotted in **Figure 1F**(IV), demonstrating a reduction in the fiber’s cross-section intensity before complete rupture. This stepwise drop in force and decrease in fiber intensity indicate that fibers fray before experiencing full breakage. This phenomenon is further illustrated in frayed fiber in **Figure 1F-II** (white arrow).

### Cyclic loading reveals permanent damage in fibrin fibers

Motivated by the dynamic nature of in vivo conditions, we next conducted cyclic loading experiments on single fibrin fibers. **Figure 2** illustrates the force-strain response of both uncross-linked and partially cross-linked fibrin fibers under cyclic loading. In this experiment, each fiber was pulled to a certain strain during loading and then released during unloading. We performed two loading cycles to discern material damage versus viscoelastic energy dissipation under the assumption that most damage would occur during the initial loading.

**Figure 2.**
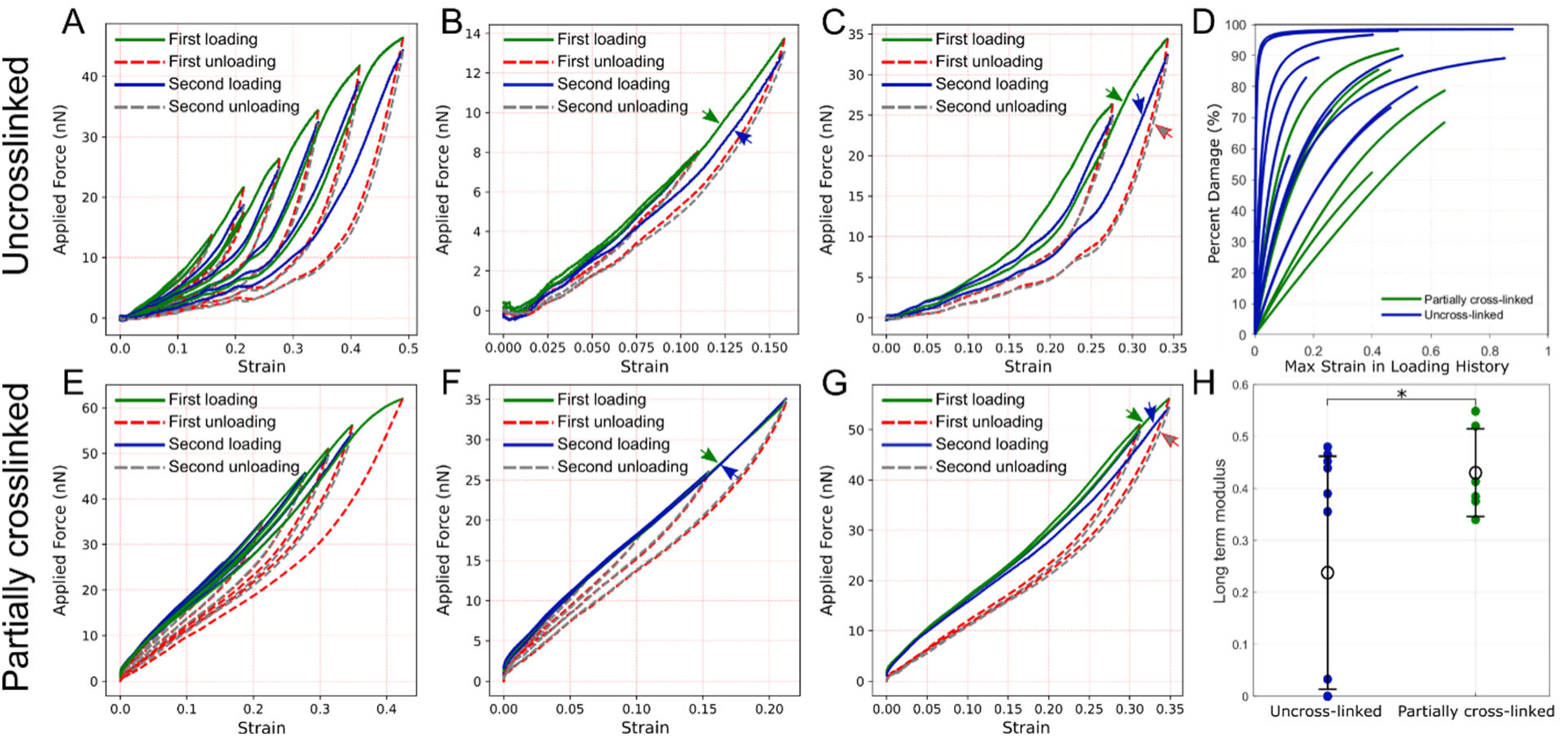
Force-strain response of single fibrin fibers in cyclic loadings. (A) the uncross-linked force-strain response in cyclic loadings. The fiber is pulled twice to each strain, named as the first and second loading/unloading. (B) the cyclic response of uncross-linked fibers in small strain (<0.20). (C) the force-strain response of the uncross-linked fiber in large strain (<0.35). Green, blue, and red/grey arrows point out to the first loading, second loading, and first/second unloading, respectively. (D) Uncross-linked fibers, shown in blue, undergo damage to a greater degree and more rapidly than partially cross-linked fibers, shown in green. (E) the force-strain response of partially crosslinked fiber in a cyclic loading. The fiber is stretched twice to each strain, named as the first and second loading/unloading. (F) the cyclic response of the fiber in (E) in small strain (<0.25). (G) the force-strain curves of the fiber in (E) in large strain (<0.35). (H) Uncross-linked fibers have a smaller long-term modulus (γ_∞_ than partially cross-linked fibers. This long-term modulus is a normalized, unitless quantity.

**Figure 2A** presents consecutive cyclic experiments for an uncross-linked fiber with increasing strain. The fiber exhibits only viscoelastic behavior at very small strains as the first and the second loadings coincide while the first loading and first unloading differ. However, permanent changes become apparent as the first and second loadings deviate from one another at ∼15% strain, illustrated in **Figure 2B**. At larger strains, shown in **Figure 2C**, the deviation between the first and second loadings becomes more pronounced, indicating more prominent damage. Interestingly, the second loading follows a different path between the first loading and the first unloading, indicating that a combination of viscoelasticity and damage occurs (discussed further in the Discussion).

**Figure 2E** depicts a similar cyclic experiment for a partially cross-linked fibrin fiber. In small strains, the first and second loading curves overlap, indicating that cross-linked fibers exhibit only viscoelasticity with no noticeable damage in small strains (up to ∼%20 strain, Fig. 2F). However, as the strain increases, the second loading again takes a different path between the first loading and unloading, suggesting permanent changes in fiber’s structure (Fig. 2G). Generally, the degree of cross-linking correlates with mechanical reversibility (the degree to which the material can recover its initial mechanical state in cyclic loadings) in fibrin fibers. As the fibrin fibers become more cross-linked, they show permanent damage in larger strains.

**Figure 2D** shows the evolution of damage calculated with a computational model of fiber deformation throughout the first loading cycle. In this figure, we plot [1 - D(W°)] ∗ 100% for ease of interpretation. The damage function, D(W°), scales down the stiffness of the fibers. In this figure, the percent damage represents the percent degradation of the original stiffness of the fiber. Uncross-linked fibers experience damage much more rapidly and to a higher degree than partially cross-linked fibers.

**Figure 2H** shows the long-term moduli calculated for all samples using the computational model. There is a significant difference between partially cross-linked and uncross-linked fibers. To perform this statistical analysis, we used Welch’s t-test in R. This modulus reflects the percentage of the material response that is purely elastic. Here, the mechanical response of partially cross-linked fibers has a larger elastic portion than uncross-linked fibers, consistent with experimental data.

Exemplary fits to the partially cross-linked and uncross-linked fibers are shown in the supplemental document (**Fig. S4-S5**). Furthermore, the parameters identified with our computational model are given for each fit (**Table S1-S2**). Interestingly, both relaxation time constants for partially cross-linked fibers are of order seconds. This is counter to the relaxation time constants for the uncross-linked fibers, where the two relaxation time constants have different orders of magnitude (**Fig. S6**). For these uncross-linked fibers, one modulus is on the order of seconds and the other is of order minutes. This means that partially cross-linked fibers could be modeled using a one-term (N = 1) Prony series, while the uncross-linked fibers need a two-term (N = 2) Prony series. Additionally, while the strain stiffening parameter, α, is higher in partially cross-linked fibers, it is not statistically significant (**Fig. S7**).

### BCARS molecular microscopy reveals the origins of molecular damage

To further probe changes in fibrin during loading with molecular insights, we used BCARS spectral imaging on fibrin gels that were cyclically loaded. As a vibrational spectroscopy technique, BCARS can reveal how different molecular modes are locally modified under different loading conditions ^24,26^. In each experiment, we cyclically deformed fibrin gels: initially pulled to ∼16% strain, then released to the original relaxed (0% strain) state, and this process was repeated a second time. Subsequently, the same gel was cyclically deformed to ∼66% strain (large strain) in a similar way. At each strain state, BCARS hyperspectral data were collected.

**Figure 3A,D** show the mean spectrum of each state for uncross-linked and partially cross-linked fibers, respectively, normalized based on the area under the curve in the Amide I region (1600 cm^−1^ to 1700 cm^−1^). Amide I region contains information about protein secondary structures, including α-helices at 1649 cm^−1^ and β-sheets at 1672 cm^−1^ ^47^. The ratio of α-helix mean intensity over the range of 1645 cm^−1^ to 1655 cm^−1^ to β-sheet mean intensity in the range of 1669 cm^−1^ to 1679 cm^−1^ is plotted in **Figure 3B** and **E** for uncross-linked and partially cross-linked fibrin gel, respectively. For uncross-linked fibers (Fig. 3B), the median of the α-helix to β-sheet ratio increases in the first small strain, followed by a further increase in the second loading. In the relaxed states (Relax1 and Relax2), the secondary structure exhibits partial reversibility even in small strains, as the ratios are reduced relative to the strained states but not entirely to the initial state. Interestingly, at large strain, the ratio increases under force to 66% strain, while the value in Relax3 is the same as Relax2. **Figure 3C** and **F** show the intensity at 3065 cm^−1^, assigned to aromatic amino acids. The amplitude of this vibration serves as an indication of water-protein interactions under force ^33^. For uncross-linked fibrin gels (Fig. 3C), there is no significant difference in this aromatic mode between loading and relaxation states at small strains. However, the median intensity in Relax1 is larger than the initial state and appears recoverable after the second loading in Relax2. At large strains, the first pull significantly reduces the 3065 cm^−1^ intensity. This intensity does not return to the same value in the second pull, signifying total, irreversible damage with this initial large deformation.

**Figure 3.**
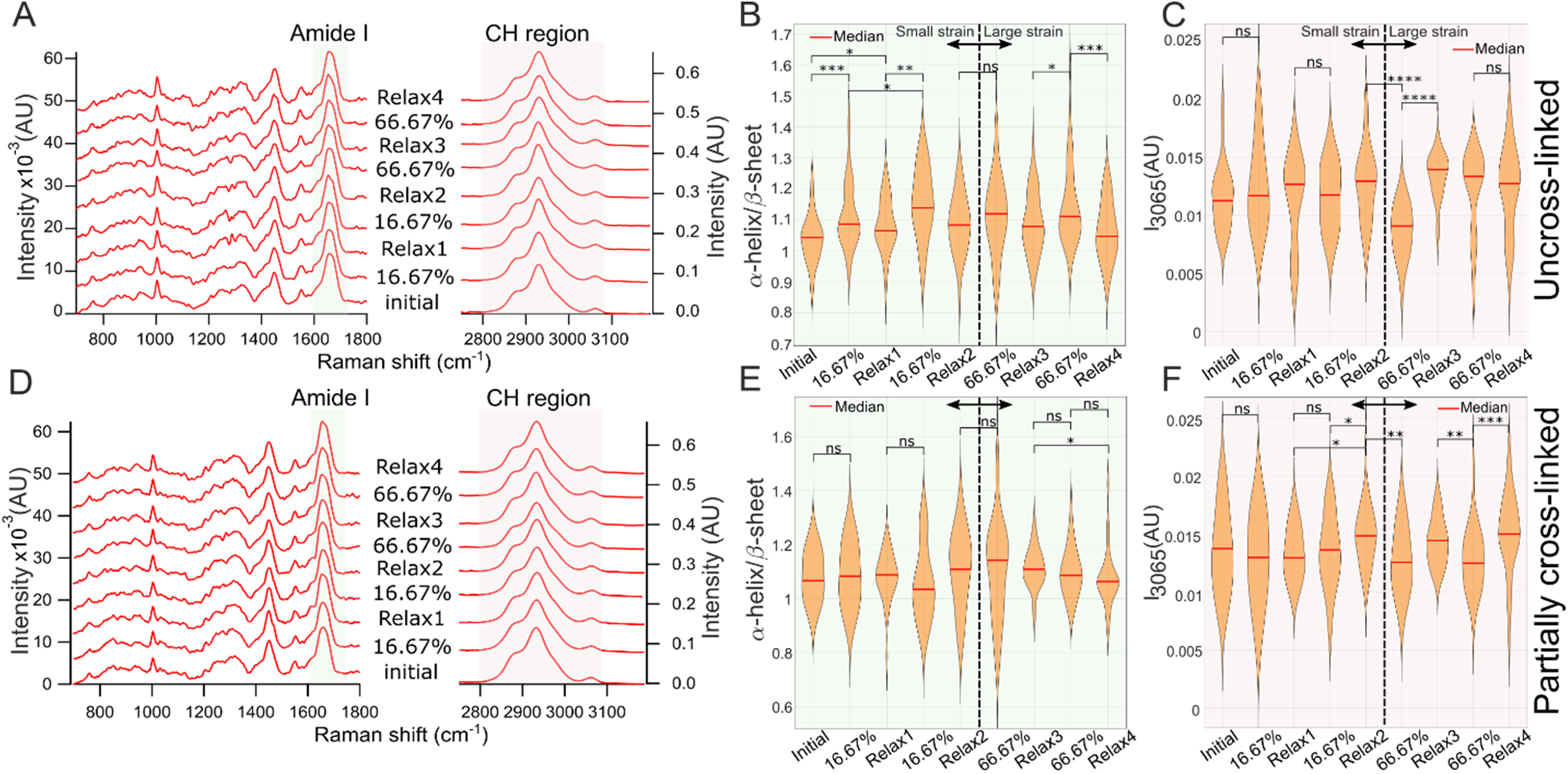
BCARS molecular spectroscopy on fibrin gel under cyclic loadings. (A) mean BCARS spectra for uncross-linked fibrin gel in different states, normalized based on area under the curve in Amide I. (B) α-helix mean intensity over β-sheet mean intensity in different states for uncross-linked fibrin fibers. (C) aromatic amino acid mean intensity at 3065 cm^−1^ for different states for uncross-linked fibers. (D) mean BCARS spectra for partially cross-linked fibrin gel in different states, normalized based on area under the curve in Amide I. (D,E) α-helix mean intensity over β-sheet mean intensity and intensity at 3065 cm^−1^ in different states of cyclic loading for partially cross-linked fibrin gel. 16.67% and 66.67% refer to the amount of strain. *, **, ***, **** show p-value smaller than 0.05, 0.01, 0.001, and 0.0001, respectively, and ns means “no significant difference.” Each state has about 30 data points from 4 and 5 experiments for uncross-linked and partially cross-linked fibrin gels, respectively.

For partially cross-linked fibers (**Fig. 3E and F**), generally smaller and less significant changes are observed at small strains compared to uncross-linked fibers. Although the BCARS data mostly do not show statistically significant differences (based on p-value ::; 0.05), in our interpretation, the observed increasing or decreasing trends may be considered reliable if they show consistency in both small or large strains or in uncross-linked fibers where larger, more pronounced changes happen. Also, we note that each state in **Figure 3A-F** is based on ∼ 30 spectra, which themselves are the mean spectra of at least 9 pixels, giving more statistical reliability to the observed trends. As shown in **Figure 3E**, the α-helix to β-sheet ratio remains almost unchanged at small strain until the second pull, where the ratio decreases sharply. This may be due to the permanent damage applied during the first loading. For cross-linked fibrin at large strain, we expected to observe more α-helix to β-sheet transition under force ^26^; however, consistent with the increase in uncross-linked fibers, the median ratio increases slightly in the first loading to 66% strain and then returns at Relax3 to almost the same value as in Relax2. Then, similar to the second pull at small strain, the α-helix to β-sheet ratio decreases in the second loading to 66% strain, followed by additional irreversible damage in Relax4, as shown by the lower value. Moving to the aromatic intensity at 3065 cm^−1^ in partially cross-linked networks (Fig. 3F), we observe that the vibration does not follow the same trend observed for uncross-linked fibers. However, the values at Relax1 and 2 are significantly different, indicating irreversible changes in the conformation of the protein, even in small strains. At larger strains, consistent with uncross-linked fibers, the aromatic amino acids’ intensity decreases in both the first and second loadings compared to the immediately preceding Relax state, with approximate reversibility observed in Relax3 and 4. The spectroscopic results from aromatic amino acids are consistent with changes in the CH_3_ intensity at 2934 cm^−1^ for different strain levels (**Fig. S8**), highlighting irreversible damage in fibrin’s tertiary structure, predominantly in uncross-linked fibers. Additionally, consistent with the results from the AFM experiments, compared to partially cross-linked fibers, large strains cause more damage (irreversible Raman intensity) to uncross-linked fibers.

### Unfolding of secondary structures reflects irreversible damage in uncross-linked fibrinogen

Different domains of a single fibrinogen molecule are shown in **Figure 4A**. A qualitative assessment of the representative structures across different pulling and relaxation states shows substantial configurational changes in γ-nodule 1 (**Fig. 4B**), while coiled-coil regions exhibit more subtle changes (**Fig. 4C and D**). In Pull 1, unfolding begins near ILE394, the pulling point in γ-nodule 1. Several secondary structure elements are completely disrupted, including: α-helices: Phe389-Thr393, Ala289-Asp291, Lys356-Ser358 and β-sheet: Lys380-Pro387. Additionally, β-sheet Ala279-Gly284 partially unfolds, splitting into two smaller β-sheets (Phe265-Val267 and Leu276-Tyr280). In Relax 1, β-sheet Ala279-Gly284 completely unfolds, while α-helix Lys356-Ser358 recovers its structure. This suggests that α-helices are more likely to recover upon relaxation. During Pull 2, α-helix Ala289-Asp291 recovers, while α-helix Lys356-Ser358 unfolds again and β-sheet Gly188-Arg197 also begins to unfold. In Relax 2, β-sheet Gly188-Arg197 recovers, while α-helix Ala289-Asp291 remains unfolded. These features in Relax 1 and Relax 2 that do not return to their original configuration represent irreversible damage.

**Figure 4.**
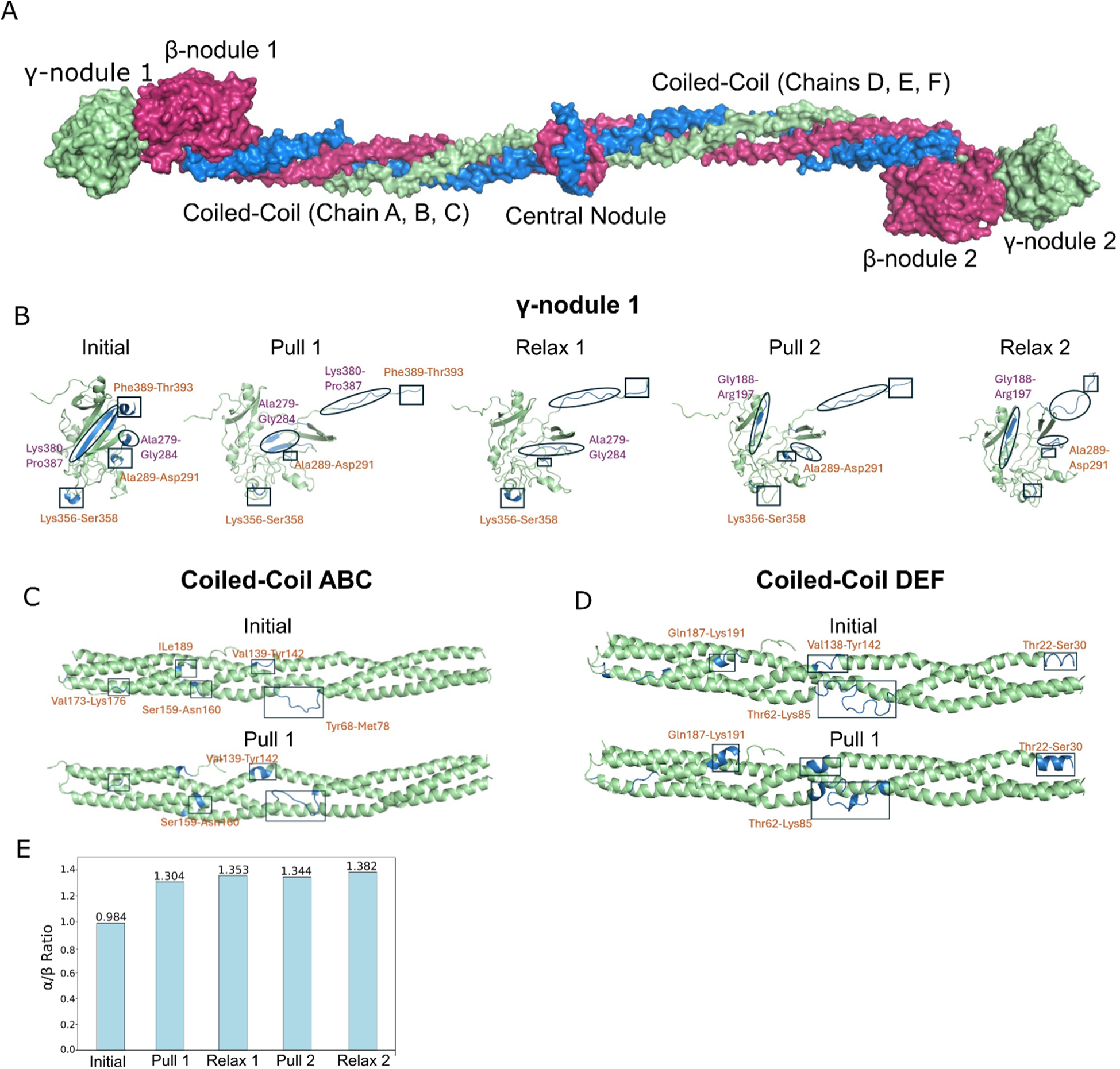
MD simulation of cyclic loading on a single molecule fibrinogen. (A) The different domains of the protein: γ-nodules 1 & 2, β-nodules 1& 2, central nodule, coiled-coil ABC, and coiled-coil DEF; (B) observed changes in γ-nodule 1 based on the selected structures representative of the experimental strain percent, changes in β-sheets represented by circle and labeled in purple, changes in α-helix represented by square and labeled in orange. Only the regions that undergo structural changes are labeled at each phase; (C) observed changes in the coiled-coil structures ABC and (D) DEF, changes in loops are represented by squares and labeled in orange (E) α-helix/ β-sheet ratio of the whole fibrinogen protein throughout cyclic loading.

Additionally, the coiled-coil structures (**Fig. 4C and D**) exhibit significant loop rearrangements throughout cyclic pulling and relaxation. During Pull 1, the following changes are observed in Coiled-Coil ABC: complete folding of loop Thr22-Ser30 and loop Val138-Tyr142, partial folding of loop Thr62-Lys85, and slight folding in the terminal loop adjacent to α-helix Gln187-Lys191. In Relax 1, the loop adjacent to α-helix Gln187-Lys191 unfolds and loop Thr62-Tyr142 forms a longer α-helix compared to Pull 1 (**Fig. S9**). Surprisingly, during Pull 2, more folding occurs in the loop adjacent to the α-helix at residue Glu172, followed by more unfolding in Relax2.

The Coiled-Coil DEF region follows a similar trend to Coiled-Coil ABC, as shown in **Figure 4D**. Several loops in Coiled-Coil DEF fold in Pull 1 including: loop Val139-Tyr142, similar to Coiled-Coil ABC, loop Ser159-Asn160, loop at ILE189, loop Val173-Lys176, and terminal loop Tyr68-Lys78. In Relax 1, no significant structural changes are observed. During Pull 2, the region Val173–Lys176 near the helix undergoes further unfolding, followed by the loop Ser159–Asn160 regaining its original structure in Relax 2 (**Fig. S10**)

The α-helix/β-sheet ratio of the fibrinogen molecule changes in response to different loading and unloading phases (**Fig. 4E**). The ratio increases in both Pull 1 and Pull 2, aligning with experimental data (**Fig. 3B**). However, during the Relax 1 and Relax 2 phases, a higher α-helix/β-sheet ratio is observed, different from experimental results. This discrepancy may be attributed to the retained strain in the relaxed structures within 6 ns of relaxation. Unlike in AFM experiments, a 0% strain state was not achieved in computational simulations.

## Discussion

### Cross-linking changes force propagation through fibrin fibers

Fibrin fibers display exceptional properties like high extensibility and non-linear viscoelastic behavior ^28,48^. Their hierarchical structure comprises aggregated, long protofibrils formed through knob-hole interactions between fibrin molecules. When force is initially applied to fibrin fibers, potentially twisted protofibrils align with the force’s direction, stretching the γ-γ cross-links between protofibrils and the αC domain—an integral component involved in lateral assembly ^19,48^. At larger strains, the fibrin molecule experiences increased force, leading to the unfolding of the helical coiled-coil and γ-nodule in the D-domain, as demonstrated by single molecule force spectroscopy and MD simulations ^17^. FXIII makes fibrin fibers more rigid as it cross-links both the αC domain and γ-nodule, increasing fibrin’s stiffness ^18,19^. The participation of cross-linking in fibrin’s response to external force results in a more solid behavior with less energy dissipation shown by our AFM experiments, as expected. Non-covalent bonds, such as hydrogen bonding, are considered sources of energy dissipation in polymers, leading to more viscoelastic behavior ^49,50^. The effect of cross-linking is also evident in small strains. Uncross-linked fibers exhibit viscoelastic behavior from small tensile strains where protofibrils align, and the unstructured αC domain is engaged as an entropic spring. Conversely, cross-linked fibrin fibers show increased elasticity and less dissipation. The decreased dissipation, particularly at low strains, underscores the presence of cross-linkers in the αC domain. Finally, independent of cross-linking, fibrin fibers underwent fraying before the final rupture, observed in both mechanical responses and fluorescence imaging. This suggests that lateral interactions between protofibrils, and therefore those involving αC domain interactions, break before the complete rupture of the protofibrils.

### Fibrin fibers show irreversible damage under load

LFM mechanical testing shows that fibrin fibers exhibit unique behavior under cyclic loading that is not pure viscoelastic, exhibiting a different behavior from the initial loading—a phenomenon known as “preconditioning” observed in various biological materials ^21,51,52^. This behavior shares similarities with Mullin’s effect, a phenomenon extensively studied in rubber-like materials ^38^. In a pure Mullin’s effect, the subsequent loading follows the previous unloading path, indicating altered material properties due to permanent damage ^53^. While a pure Mullin’s effect entails irreversible damage, which leads to different loading/unloading paths, viscoelastic materials dissipate some energy, leading to a more substantial decay in the unloading curve. Although the damage from the Mullin’s effect is largely irrecoverable, materials can recover some energy loss due to viscoelasticity. For a pure viscoelastic material, the second loading would follow the first loading with a Lissajous loop observable during unloading. In fibrin fibers, we find the new loading deviates from both the prior loading and unloading paths, instead following a new trajectory between these curves, a result of a combined Mullin’s-like effect and viscoelasticity (**Fig. 2C,G**), similar to the mechanical behavior observed in electrospun poly(ε-caprolactone) single fibers ^54^. In small strains, fibrin fibers primarily exhibit viscoelastic behavior, where the first and second loadings coincide. This trend persists up to larger strains (∼ 20%) in partially cross-linked fibers, after which permanent damage is seen; however, uncross-linked fibers start showing permanent damage at ∼15% strain. Cross-linking and covalent bond formation delay structural damage in fibrin fibers up to larger strains.

### Fibrin’s unfolding pattern depends on its loading history

The BCARS results complement the LFM mechanical data, providing insights into the molecular structural alterations occurring within fibrin fibers. In uncross-linked fibers, a significant increase in the α-helix to β-sheet ratio is observed under a small strain of 16%, a phenomenon notably absent in partially cross-linked fibers where the ratio remains relatively stable throughout the first and second loadings. This significant unfolding apparently results in more permanent changes in fibrin’s secondary structure as the ratio does not return to the original value that it was in the initial state. This signifies only partial refolding of the chains, even in response to small strains.

The unexpected increase in the α-helix to β-sheet ratio during cyclic loadings in uncross-linked fibers contrasts with the typically observed transition towards a decreasing ratio under load ^25,26^. This may be explained by MD simulations for uncross-linked fibrinogen molecule showing that both α-helices and β-sheets unfold under the applied force in both γ-nodule and coiled-coil structures. While some α-helices are able to recover back to their previous state, β-sheets show minimal recovery. Even larger β-sheets, such as Gly188-Arg197, which are located away from the pulling point (ILE394), undergo unfolding. This suggests that β-sheets are more susceptible to damage compared to α-helices at least at low strains. Interestingly, in the coiled-coil regions, loops play a crucial role in the folding and unfolding dynamics of the entire structure. The loss of loops during the initial pulling phase may also explain the increased α-helix/β-sheet ratio.

Conversely, in partially cross-linked fibers, Raman spectroscopy showed no significant difference among various states, which exhibit less force propagation to the fibrin molecules and, hence, fewer changes in their secondary structure under small or even large strains (up to 66%). This is consistent with mechanical results of partially cross-linked fibers where less damage is observed in small strains. Also, cross-linked fibrin fibers show permanent damage in larger strains compared to uncross-linked fibers ^19^, meaning cross-linking reduces the applied force to the fibrin molecules and, hence, significant molecular changes. Additionally, the α-helix to β-sheet ratio differs in two consecutive loadings with the same amount of strain. In uncross-linked fibers, compared to the first pull, the ratio is larger in the second pull, but in partially cross-linked fibers, it becomes smaller. We attribute this difference to the damage that is applied to different domains in the fibrin molecule. The stiffness in protein’s different domains changes upon the first loading, which causes a redistribution of deformation and hence a different mechanical response in the second loading (with the same amount of strain).

### 3D conformation of fibrin changes under load

Aromatic amino acids, mainly present in the D-domain of the fibrinogen molecule (depicted in **Table S3**), exhibit hydrophobic properties with a distinct peak at ∼3065 cm^−1^ ^33,56^ and play a role in the tertiary (3D) conformation of fibrin. This has led to a decrease in the Raman intensity at 3065 cm^−1^, illustrated in large strains in Figure 3C**, E**. While this pattern is observed in uncross-linked fibrin fibers, partially cross-linked fibers exhibit this behavior only in large strains. We attribute this difference to the effect of cross-linking that limits force propagation through fibrin’s molecule until large strains are reached.

Additionally, aromatic amino acid intensity demonstrates significant irreversible changes in large strains for uncross-linked fibers, highlighting 3D conformational changes in the tertiary structure of the protein, likely coming from the D-domain as it has the largest portion of aromatic amino acids (**Table S3**). However, more reversibility is observed for the same level of strain in cross-linked fibers, consistent with mechanical testing results.

## Conclusion

Fibrin fibers, with their hierarchical structure and unique mechanical properties, are integral to the clotting process. Their behavior under dynamic forces diverges from their response to static loading, highlighting the complexity of their mechanical response. Permanent changes in secondary structure and 3D conformation, influenced by loading history, significantly impact fibrin’s mechanical behavior. This understanding enriches our insight into clot behavior under dynamic blood flow pressure or muscle contractions, offering valuable perspectives for understanding the physiological processes and potential therapeutic interventions in clot-related pathologies.

## Acknowledgements

We acknowledge support from the National Science Foundation through grants 2105175, 2235856, 2127925, and 2046148, the Welch Foundation through grants F-2008-20220331, F-2120, the National Institutes of Health through grant R01GM106137, and the Office of Naval Research through grant N00014-23-1-2575.

## Author contributions

S.N. and S.H.P. conceived the study. S.N. performed all experiments in the study with help from C.M.J. and D.W. M.J.L. and M.K.R. performed the modeling of cyclic loading and damage at the fiber level. M.M. and P.R. performed the molecular dynamics simulations of fibrin protein structure under cyclic loading. S.N. and S.H.P. wrote the manuscript with input from all authors. S.H.P. supervised the study.

