## Supporting figures and tables for "Loading causes molecular damage in fibrin fibers"

Table of Contents:

Figures S1 – S10

Tables S1 – S3

**Supporting Figures**

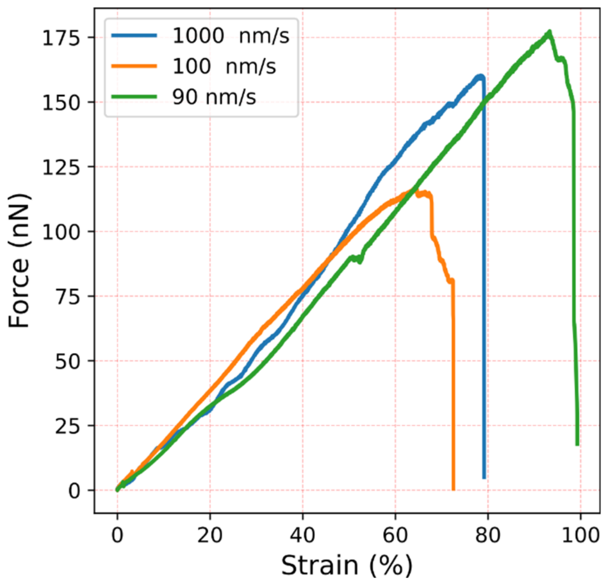

***Figure S1.*** *Force-strain curves for three single fibrin fibers with different loading rates.*

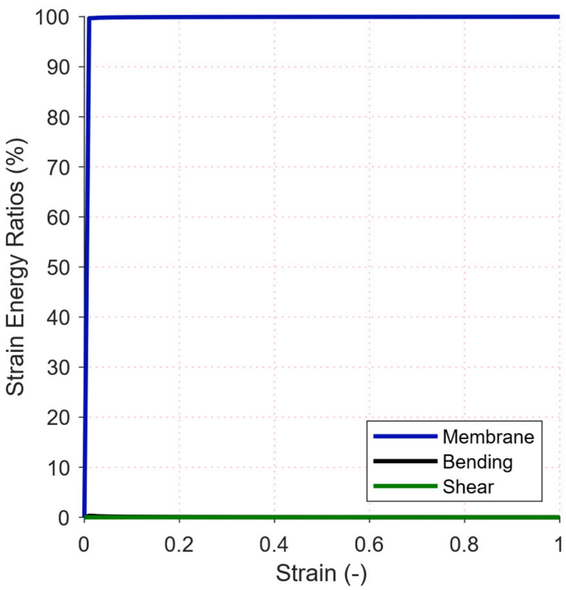

***Figure S2.*** *Strain energy contributions to a fiber under lateral deflection. This was calculated from a finite element model of a fiber under the same loading scheme as in the atomic force microscopy experiments. Membrane energy accounts for almost all of the strain energy of the system, meaning that fiber stretching plays the predominant role at strains relevant for these experiments. Because of this, we exclude fiber bending and shearing in our computational model.*

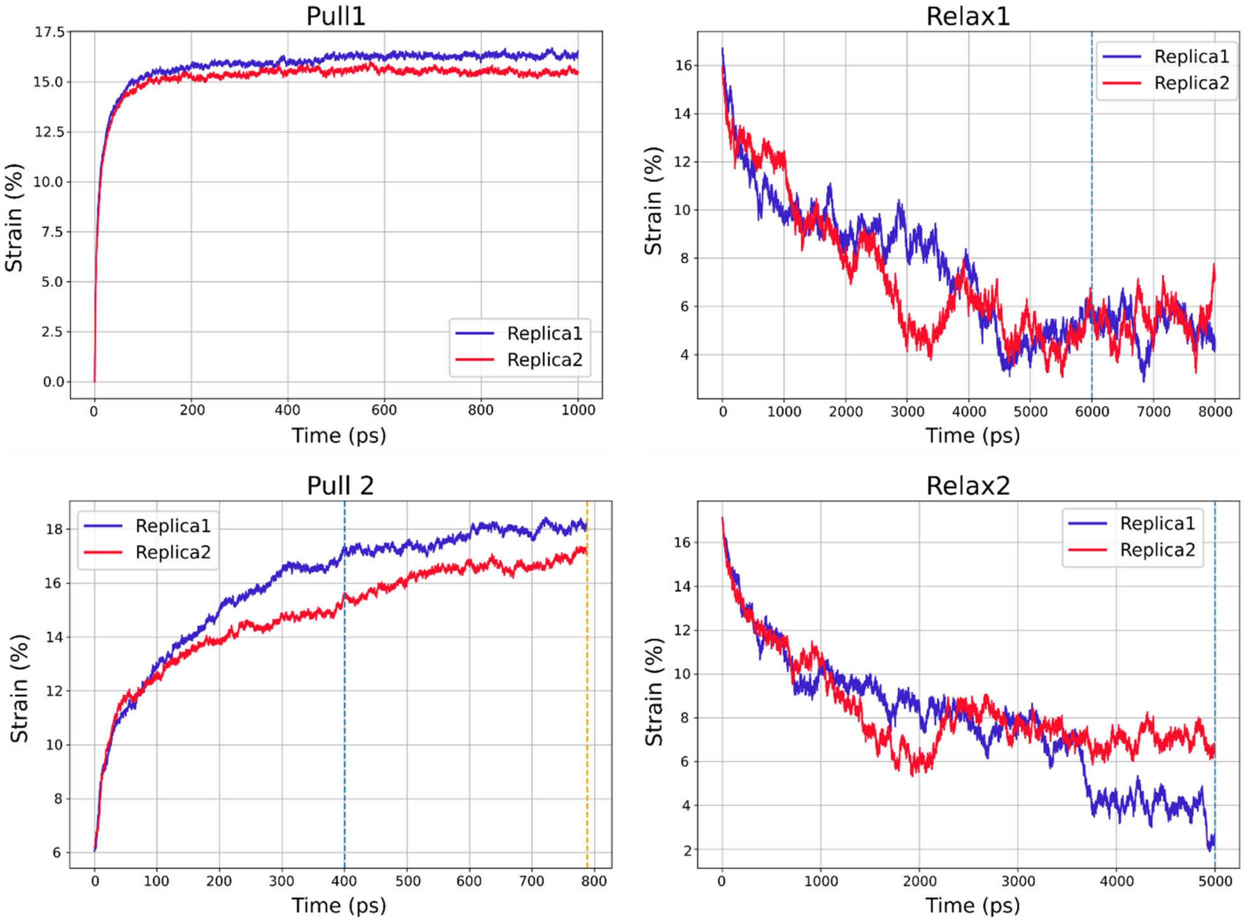

***Figure S3***. *Strain vs Time for the cyclic pulling and relaxation phases (Pull1, Relax1, Pull2, and Relax2). Dashed lines represent the selected frames utilized as representative structures of each phase. The last frame of 1ns in Pull1 was used as representative structure and the last frame of 6ns in Relax1 was used as representative structure for the two independent replicas. For Replica 1 of Pull 2, the last frame at 400ps was used as representative structure, while the last frame of 800ps was used as representative structure for Replica 2 of Pull2. The last frame of 5 ns relaxation time in Relax 2 was used as representative structure for the two replicas.*

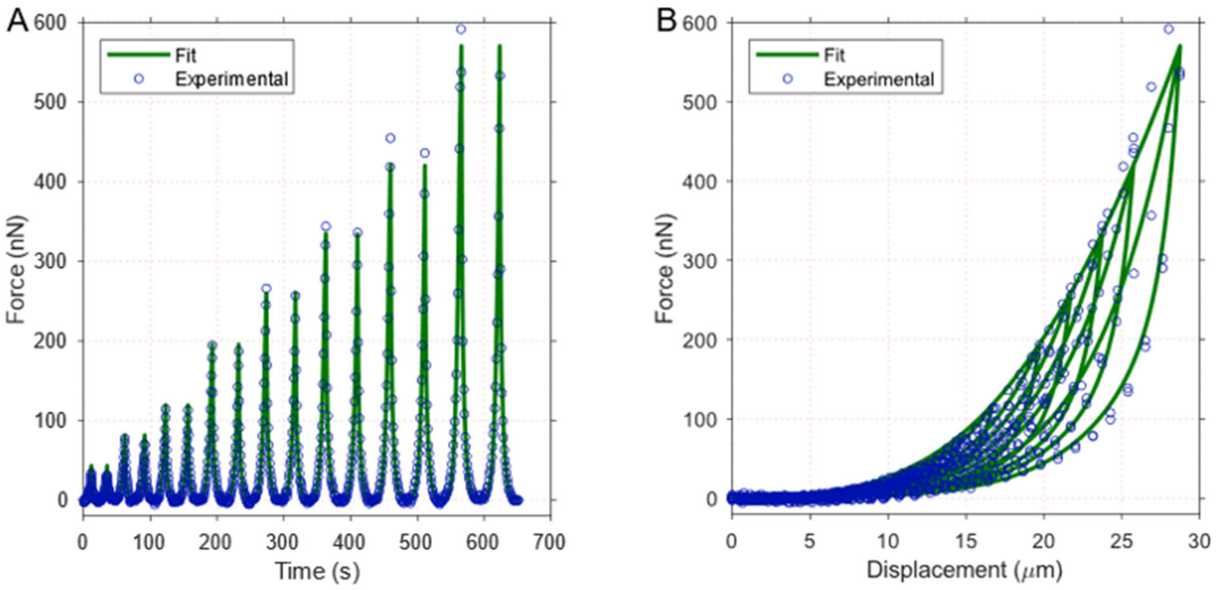

***Figure S4.*** *Exemplary fit to experimental data for cross-linked fibers: A) force-time curve, B) force-displacement curve. NMSE = 0.9929.*

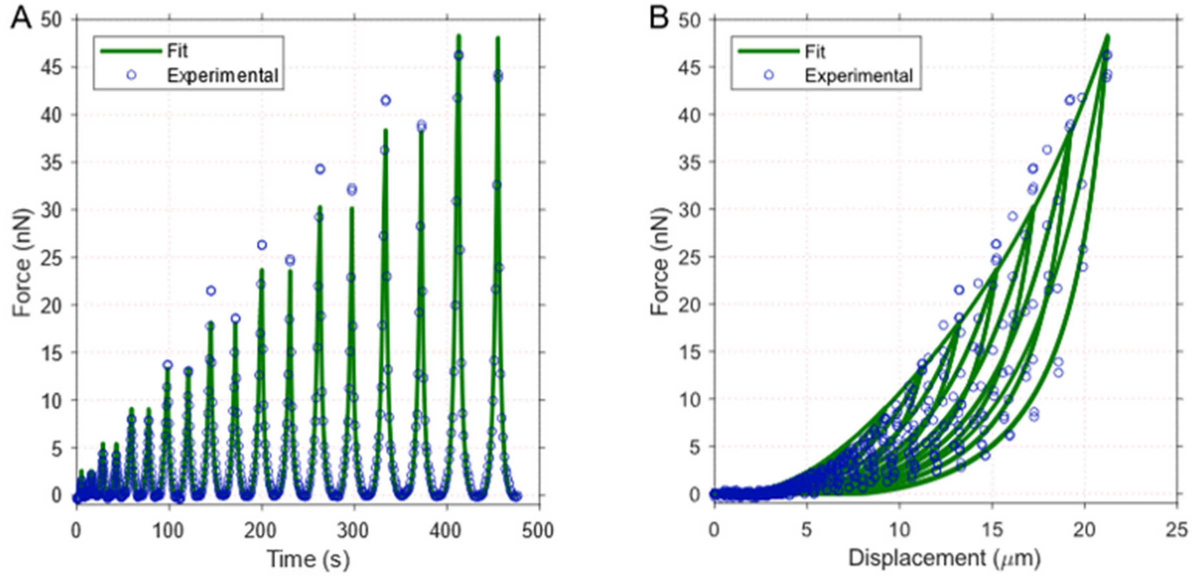

***Figure S5.*** *Exemplary fit to experimental data for cross-linked fibers: A) force-time curve, B) force-displacement curve. NMSE = 0.9876.*

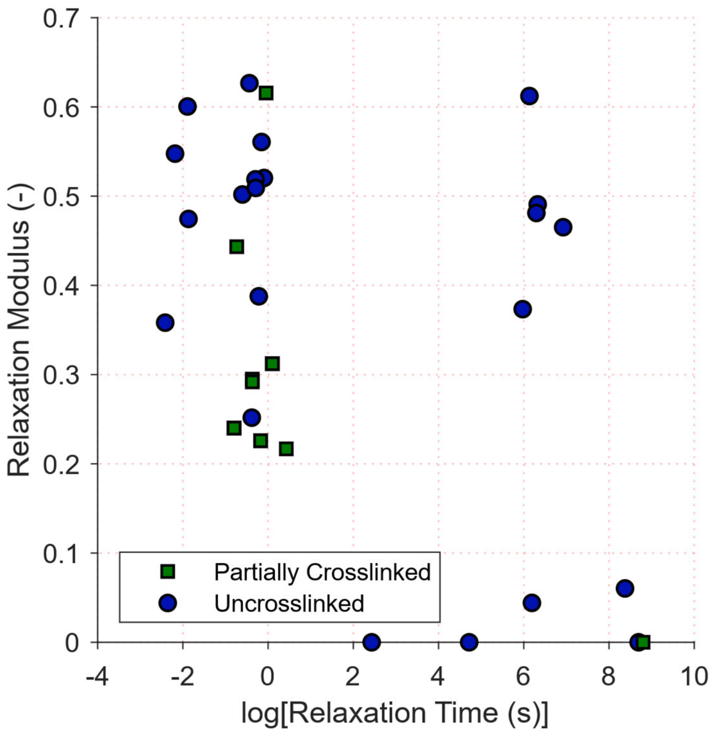

***Figure S6.*** *Relaxation times and their respective modulus. The partially cross-linked fibers have non-unique relaxation times, whereas uncross-linked fibers have two unique relaxation times. Relaxation times for partially cross-linked fibers are on the order of seconds, while relaxation times for uncross-linked fibers are on the order of seconds and minutes.*

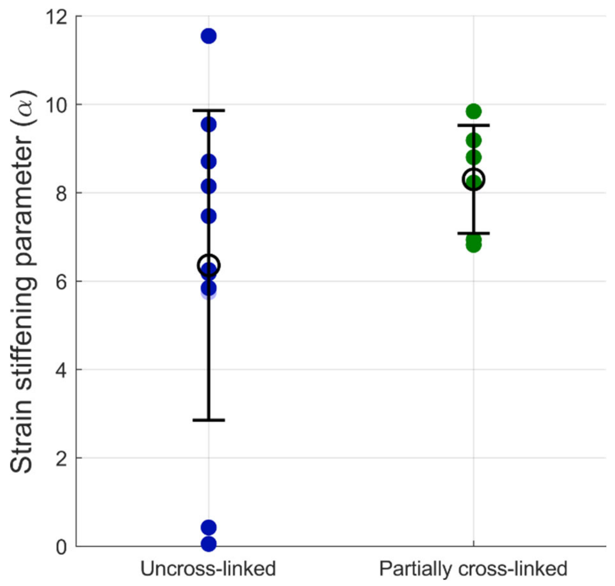

***Figure S7.*** *The partially cross-linked fibers have higher strain stiffening parameters than their uncross-linked counterparts, but it is not statistically significant.*

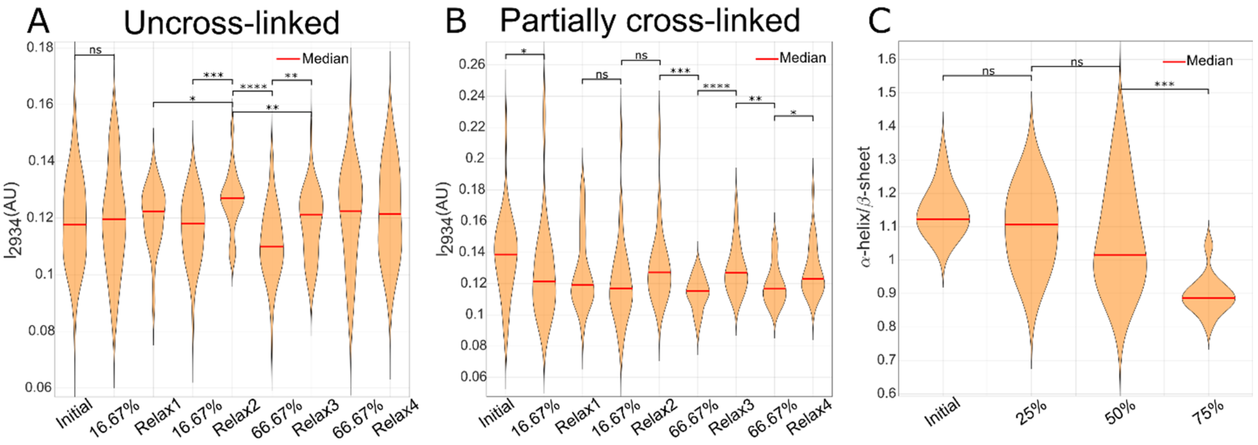

***Figure S8.*** *$CH_3$ peak mean intensity at 3065 $cm^{-1}$ for different cyclic states for (A) uncross-linked fibers and (B) partially cross-linked fibrin gels. (C) $\alpha$-helix mean intensity over $\beta$-sheet mean intensity in acyclic loading for partially cross-linked fibrin fibers. The numbers show the strain. *, **, ***, **** show p-value smaller than 0.05, 0.01, 0.001, and 0.0001, respectively. Each state in (A) and (B) consists of about 30 data points from 4 and 5 experiments for uncross-linked and partially cross-linked fibrin gels, respectively. Each state in (C) includes about ten samples from one acyclic experiment.*

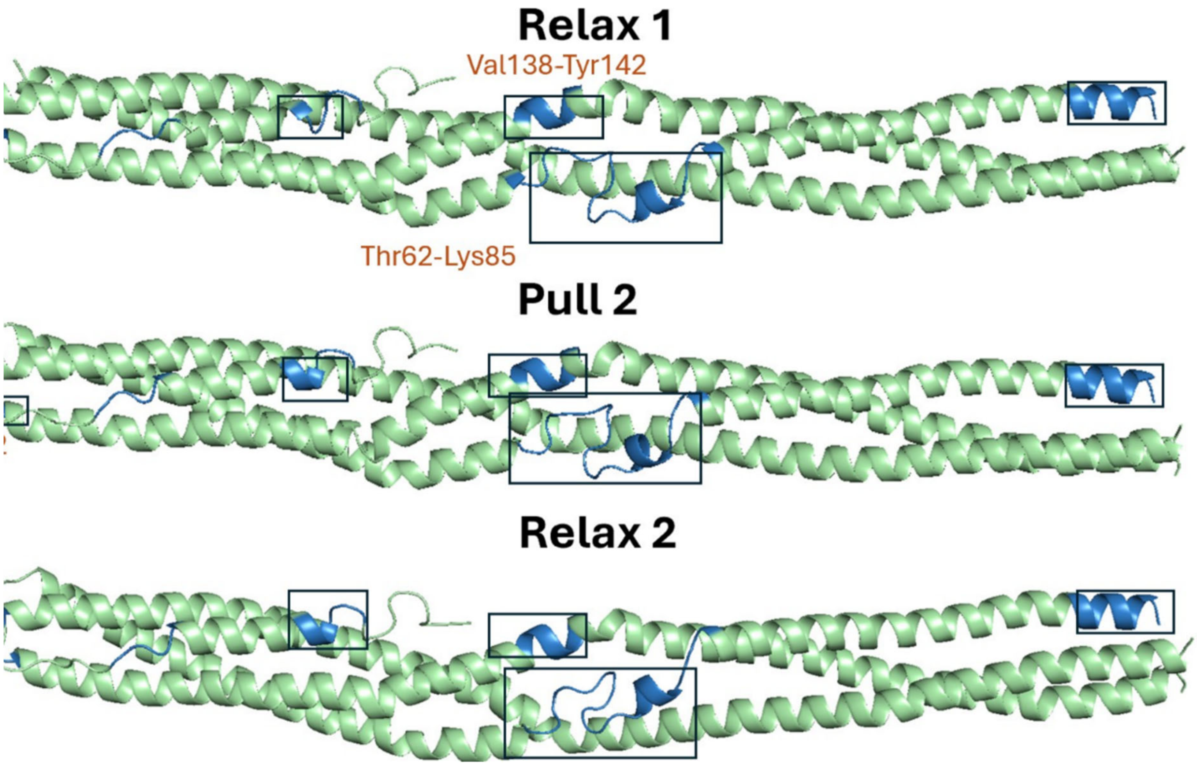

***Figure S9****. The structural changes of the coiled-coil ABC after pull 1. The changes in the loop and helices represented by boxes and the changing parts in each phase are labeled in orange.*

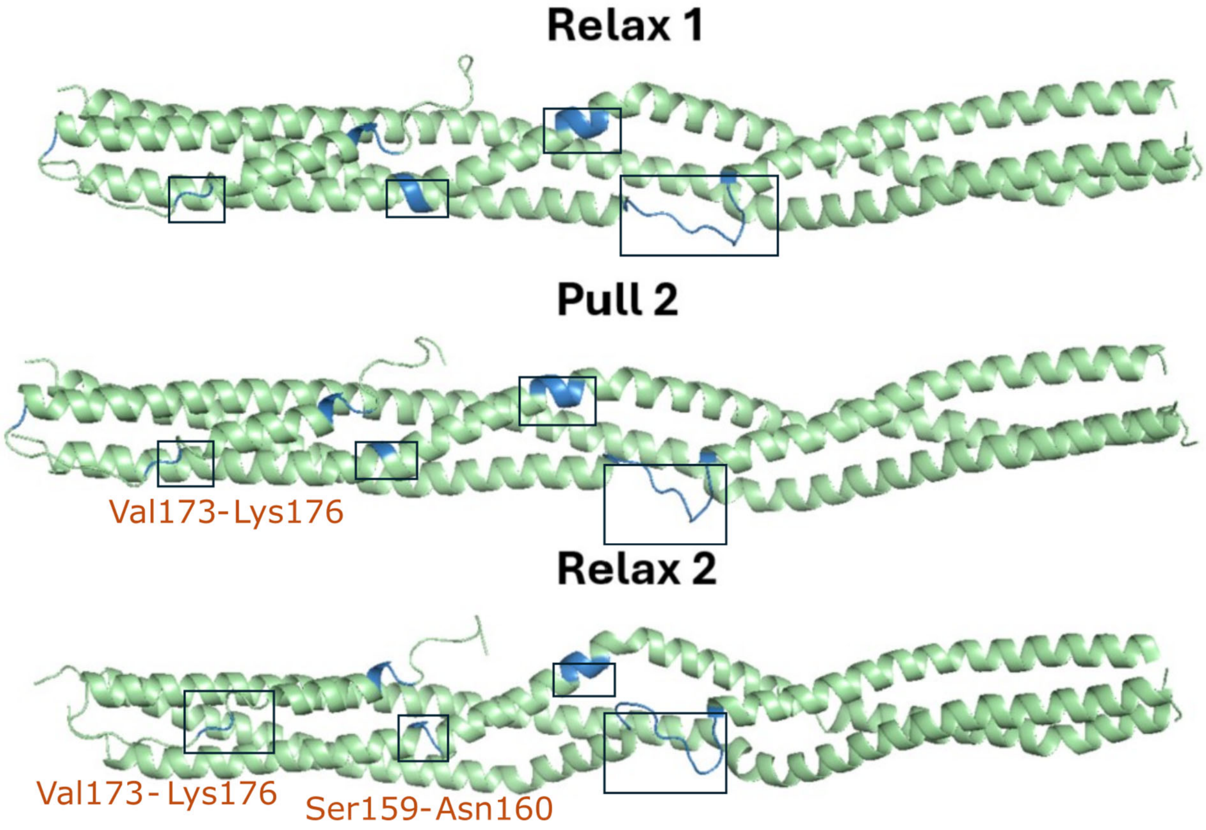

***Figure S10***. *The structural changes of the coiled-coil DEF after pull 1. The changes in the loop and helices represented by boxes and the changing parts in each phase are labeled in orange.*

### Supporting Tables

***Table S1.*** *Parameters for partially cross-linked samples. Note that, because we did not have accurate measures of diameter, the values for $\mu$ and $d_2$ are only valuable for comparisons between these samples.*

| $\mu\ (MPa)$ | $\alpha\ (-)$ | $d_1(-)$ | $d_2(MPa^{-1/2})$ | $\gamma_1(-)$ | $\gamma_2(-)$ | $\tau_1(-)$ | $\tau_2(-)$ | NMSE |
|---|---|---|---|---|---|---|---|---|
| 100 | 9.8 | 4e-4 | 4.5 | 0.22 | 0.44 | 1.5 | 1 | 0.99 |
| 20 | 8.8 | 9e-10 | 2.6 | 0.62 | 0.00 | 1.0 | 6600 | 0.97 |
| 30 | 6.8 | 9e-9 | 2.7 | 0.31 | 0.31 | 1.1 | 1 | 0.97 |
| 30 | 8.2 | 3e-9 | 0.9 | 0.24 | 0.24 | 0.4 | 1 | 0.92 |
| 80 | 6.9 | 6e-12 | 0.7 | 0.30 | 0.29 | 0.7 | 1 | 0.97 |
| 20 | 9.2 | 3e-9 | 0.7 | 0.23 | 0.23 | 0.8 | 1 | 0.96 |

***Table S2.*** *Parameters for uncross-linked samples. Note that, because we did not have accurate measures of diameter, the values for $\mu$ and $d_2$ are only valuable for comparisons between these samples.*

| $\mu\ (MPa)$ | $\alpha\ (-)$ | $d_1(-)$ | $d_2(MPa^{-1/2})$ | $\gamma_1(-)$ | $\gamma_2(-)$ | $\tau_1(-)$ | $\tau_2(-)$ | NMSE |
|---|---|---|---|---|---|---|---|---|
| 9200 | 5.7 | 0.01 | 0.2 | 0.5 | 0.1 | 0.2 | 4000 | 0.96 |
| 30 | 6.2 | 0.04 | 0.7 | 0.5 | 0.0 | 0.9 | 100 | 0.97 |
| 30 | 7.5 | 4e-12 | 1.6 | 0.6 | 0.0 | 0.9 | 10 | 0.98 |
| 40 | 8.1 | 9e-6 | 1.8 | 0.5 | 0.5 | 0.5 | 1000 | 0.91 |
| 40 | 6.2 | 8e-12 | 0.8 | 0.6 | 0.4 | 0.6 | 400 | 0.92 |
| 900 | 0.1 | 3e-5 | 0.6 | 0.4 | 0.3 | 0.1 | 1 | 0.99 |
| 40 | 9.5 | 7e-10 | 0.9 | 0.5 | 0.5 | 0.7 | 600 | 0.97 |
| 60 | 8.7 | 7e-11 | 0.8 | 0.5 | 0.5 | 0.7 | 500 | 0.94 |
| 300 | 0.4 | 0.03 | 0.5 | 0.6 | 0.1 | 0.2 | 500 | 0.98 |
| 90 | 11.5 | 9e-10 | 0.6 | 0.4 | 0.6 | 0.8 | 500 | 0.97 |
| 600 | 5.8 | 0.02 | 0.1 | 0.5 | 0.0 | 0.1 | 6000 | 0.99 |

In the above tables, NMSE is defined as follows

$$NMSE = 1 - \frac{\sum_i^N (F_i^{exp} - F_i^{num})^2}{\sum_i^N (F_i^{exp} - mean(F^{exp}))^2} \tag{1}$$

***Table S3.*** *The number of aromatic amino acids (highlighted in green) and amino acids with $CH_3$ groups (highlighted in orange) in different domains of fibrinogen molecule. The number was obtained from 3GHG crystal structure [1], except the numbers in the $\alpha$C-domain, which were counted based on the $\alpha$ chain sequence in [2].*

| Amino acid | E-domain | Coiled-coil | D-domain | $\alpha$C-domain |
|---|---|---|---|---|
| Phe | 4 | 10 | 42 | 26 |
| Tyr | 4 | 26 | 56 | 10 |
| Trp | 4 | 2 | 44 | 16 |
| Sum | 12 | 38 | 142 | 52 |
| Percent | %4.9 | %15.6 | %58.2 | %21.3 |
| Leu | 6 | 84 | 42 | 18 |
| Met | 0 | 16 | 36 | 6 |
| Ile | 0 | 52 | 48 | 10 |
| Ala | 8 | 28 | 52 | 28 |
| Val | 2 | 50 | 40 | 26 |
| Thr | 6 | 28 | 70 | 86 |
| $\#CH_3$ | 30 | 444 | 418 | 228 |
| Percent | %2.7 | %39.6 | %37.3 | %20.4 |
